## Supplemental File -- Raw Blots for "A Soluble Platelet-Derived Growth Factor Receptor-β Originates via Pre-mRNA Splicing in the Healthy Brain and is Differentially Regulated during Hypoxia and Aging"

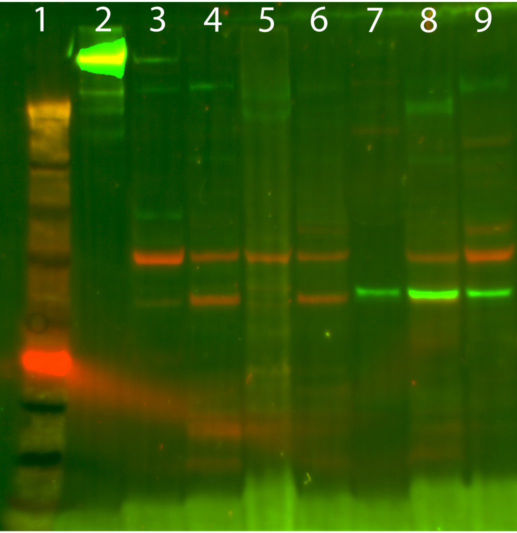


PDGFRb

GAPDH

1 - Marker
2 - Recombinant PDGFRb
3 - Skelatal Muscle
4 - Heart
5 - Intestine
6 - Liver
7 - Serum
8 - Kidney
9 - Brain

in support of Figure 1

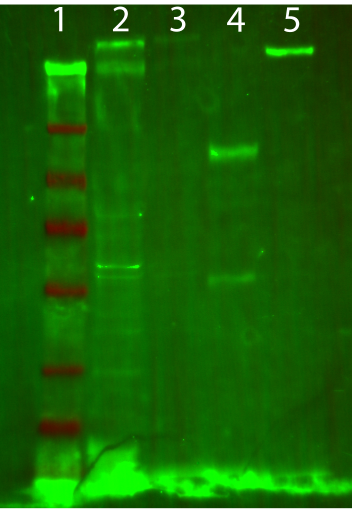


1 - Marker
2 - Whole Brain Lysate
3 - Depleted Lysate
4 - Co-IP Sample
5 - Recombinant PDGFRb

**PDGFRb**

in support of Fig. 3

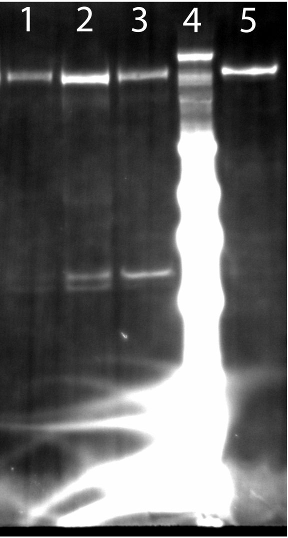


PDGFRb
1 - P7 Brain
2 - P21 Brain
3 - P90 Brain
4 - Marker
5 - Recombinant PDGFRb

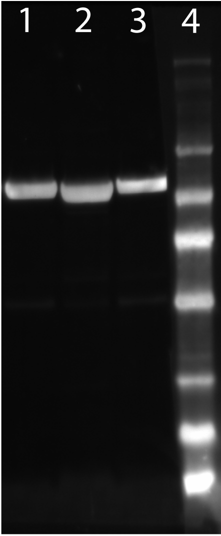


Alpha-tubulin
1 - P7 Brain
2 - P21 Brain
3 - P90 Brain
4 - Marker

in support of Figure 6
